# Early life adversity shapes adult behavior in a free-ranging primate

**DOI:** 10.64898/2026.07.21.739374

**Authors:** Sam K. Patterson, Josué E. Negron-Del Valle, Daniel Phillips, Angelina Ruiz-Lambides, Cayo Biobank Research Unit, Noah Snyder-Mackler, Lauren J.N. Brent, James P. Higham

**Affiliations:** Department of Anthropology, University of Notre Dame; Notre Dame, IN, USA; School of Life Sciences, Arizona State University; Tempe, AZ, USA; Caribbean Primate Research Center, University of Puerto Rico; San Juan, Puerto Rico; Department of Neuroscience, University of Pennsylvania, Philadelphia, PA, USA; Center for Evolution and Medicine, Arizona State University; Tempe, AZ, USA; School for Human Evolution and Social Change, Arizona State University; Tempe, AZ, USA; Centre for Research in Animal Behaviour, University of Exeter; Exeter, United Kingdom; Department of Anthropology, New York University; New York, NY, USA; New York Consortium in Evolutionary Primatology; New York, NY, USA

**Keywords:** behavior, sociality, development, early life adversity, primates

## Abstract

Early life adversity (ELA) causes lasting psychological and behavioral consequences in humans, with parallel stress and fear responses observed in captive animals. How naturally occurring ELA shapes behavior in non-captive animals remains poorly understood. Investigating these effects across socioecological environments is important for identifying the evolutionary pressures shaping early life sensitivities, and determining whether these behavioral syndromes have deep evolutionary roots. Here, we tested the effects of eight forms of ELA on adult behavior using a large dataset (N=313 males, 346 females) from a free-ranging population of rhesus macaques (*Macaca mulatta*). We measured adult behaviors indicative of stress and the tenor of social interactions, including agonism, vigilance, self-directed behaviors, and affiliation. Individuals exposed to more ELA received more aggression and less grooming, were more submissive, and were less vigilant. While high status buffered the effects of ELA on grooming, it amplified effects on aggression. Our findings demonstrate that in a large sample of a free-ranging primate, early life environments impact behavior into adulthood. While some behaviors are consistent with patterns in humans and captive animals, suggesting an evolutionarily conserved ELA syndrome, other findings add new insights into the various ways individuals interact with their environment as a function of ELA.

## Introduction

Children exposed to adversity are at higher risk of developing mental health illnesses and risky behavioral tendencies. In the US, adults who experienced adversity during childhood (i.e. early life adversity) showed a prevalence of anxiety disorders twice as high as expected (Whitaker et al., 2021), a finding echoed by WHO data linking early life adversity to mood, anxiety, behavior, and substance disorders (Kessler et al., 2010). Early life adversity is associated with behaviors like a higher probability of smoking cigarettes, drinking alcohol, poor diet, impulsivity, and violence (Lovallo, 2013; Salo et al., 2022; Terrell et al., 2019; Veenema, 2009; Whitaker et al., 2021), which could contribute to the increased susceptibility to diseases, poorer physical health, and reduced life expectancy that has been observed in association with early life adversity. Loneliness, an important dimension of mental and physical health, is also more prevalent among older adults who experienced early life adversity (Furuya & Wang, 2023).

While a large body of empirical work supports the connection between early life adversity and later life outcomes, progress lags due to considerable sociocultural and logistical challenges. First, different forms of adversity, such as limited access to health care and limited access to quality foods, tend to co-occur and persist into adulthood, limiting the ability to isolate the effects of environmental experiences at specific developmental time points (Mersky et al., 2017). Second, because humans are long-lived, it is challenging to collect the longitudinal data across decades to directly assess experiences in early life and outcomes in adulthood (Snyder-Mackler et al., 2020). Even when sufficient long-term research infrastructure is in place, it can be hard to retain people in studies over time (Bhamra et al., 2008; Heid et al., 2018).

Nonhuman animal models can help overcome these hurdles involved with human studies. Animals live in social systems with fewer socio-cultural complexities, and as a result, different environmental adversities do not tend to co-occur or persist across life stages, allowing researchers to disentangle and isolate the effects of different forms of adversity at specific times (Evans & Kim, 2010; Snyder-Mackler et al., 2020). Further, most animals live shorter lives than humans, making it possible to track animals over relatively longer periods of their lives (Austad & Fischer, 1992; Bronikowski et al., 2011; Chiou et al., 2020; Emery Thompson et al., 2020), including from birth till death (Hayes et al., 2017; Kappeler et al., 2012; Kerth, 2022; Mann & Karniski, 2017; Sheldon et al., 2022; Smith et al., 2017; Tung et al., 2010).

Experimental research with captive animals has demonstrated how early life adversity alters neurobiology and behavior (Dettmer & Suomi, 2014; Harlow & Zimmermann, 1959; Suomi et al., 2011; Whitehouse et al., 2017). Stress and anxiety are typically operationalized through high rates of repetitive or self-directed behaviors (e.g., pacing, rocking, self-scratching, self-grooming) (Dettmer et al., 2012; Mikheenko et al., 2015; Raper et al., 2018; Whitehouse et al., 2017), and fear, impulsivity, and reactivity are measured via high levels of submissive and aggressive behaviors (Barros et al., 2000; Barros & Tomaz, 2002). Rodents and macaques are frequently used as model organisms for these studies. When experimentally exposed to early life adversity such as separation from their mothers, rodents and macaques typically display high rates of fear, anxiety-like behavior, impulsivity, aggression, and social behavioral deficiencies (Bonapersona et al., 2019; Condon et al., 2022; Haller et al., 2014; Kraemer, 1992; Pryce et al., 2005; Rosenblum & Andrews, 1994; Suomi, 1997; Veenema, 2009).

However, experimental studies on other species remain relatively scarce and yield conflicting results. Honey bees (*Apis mellifera*) reared in high-aggression environments exhibit high rates of aggression as adults (Rittschof et al., 2015); whereas field crickets (*Gryllus integer*) exposed to cues of high competition become less aggressive and less likely to be dominant in adulthood (DiRienzo et al., 2012). Cichlids (*Neolamprologus pulcher*) treated with cortisol during early life show heightened aggression during social challenges, resulting in longer-lasting conflict, greater energetic expenditure, and poorer overall social performance (Reyes-Contreras et al., 2019).

While captive animal models show that early life adversity can produce behavioral outcomes potentially comparable to human anxiety, reactivity, and loneliness, we do not know if similar patterns arise under natural conditions. Captive environments lack the natural socioecological pressures that shape behavioral strategies, leaving a gap in our understanding. How does naturally occurring adversity shape behavioral phenotypes including self-directed, agonistic, and affiliative behaviors in naturally varying social and ecological contexts?

There are a small but growing number of studies investigating the ways in which early life adversity predicts adult behavior and health in natural animal populations. To date, behavioral studies have focused on affiliative behaviors and social connectedness in primates. Wild female baboons that experience early life adversity grow up to be more socially isolated than baboons without early life adversity, and this isolation is at least partially due to attracting less affiliative social attention from group-mates (Lange et al., 2023; Patterson et al., 2022; Rosenbaum et al., 2020; Tung et al., 2016). Social isolation is associated with poorer health outcomes and reduced survival across a range of social species (Snyder-Mackler et al., 2020). Early life adversity can also influence interaction style, or the ways in which individuals interact with others and their environment (Patterson et al., 2022), which in turn could shape their social relationships and health. Studying a suite of behaviors, including and in addition to affiliative interactions, can help contextualize how individuals navigate their environment and the role this may play in shaping their social connections, health, and evolutionary fitness.

Here, we study early life adversity and adult behavior in a free-ranging population of rhesus macaques on Cayo Santiago, Puerto Rico. Rhesus macaques live in multi-male, multi-female groups and form differentiated social relationships and dominance hierarchies. Infant macaques face similar early life adversities as humans such as parental loss, limited access to food, and natural disasters. Among the macaques on Cayo Santiago, exposure to early life adversity reduces life expectancy (Gonzalez et al., 2023; Patterson et al., 2024). The survival effects of early life adversity may be mediated or moderated by numerous factors such as social dominance rank, social relationships, behavior, and/or health. We investigate the link between early life adversity and behaviors. We predict that early life adversity will be associated with higher levels of self-directed behavior, vigilance, aggression, and submissive behaviors because all of these behaviors can be indicative of stress or more reactive phenotypes. We predict that macaques with early life adversity will receive higher rates of aggression from others, possibly because they are easy physical targets or because they may be more aggressive and unpredictable themselves. Given previous links between early life adversity and lower social connectivity in wild baboons, we predict that macaques with early life adversity will exhibit lower rates of approaching and grooming others and lower rates of being approached and groomed by others. Furthermore, characteristics like sex and dominance rank could shape the relationship between early life adversity and behavior. On average, male rhesus macaques experience more energetically costly developmental trajectories than females, so males are predicted to be more vulnerable to adversities like nutritional constraints (Kulik et al., 2015; Schwartz & Kemnitz, 1992; Turcotte et al., 2022). Additionally, given sex-related variation in social relationships and hierarchies (Blomquist et al., 2011; Higham & Maestripieri, 2014; Kulik et al., 2015; Weiß et al., 2016), we might expect males and females to employ different behavioral strategies as a function of early life adversity. Finally, we predict high social status in adulthood will mitigate the effects of early life adversity on these behavioral outcomes such that high ranking individuals exposed to adversity will show lower levels of self-directed behaviors, vigilance, and agonism but higher levels of affiliative interactions than low ranking individuals exposed to adversity.

## Methods

### Study system

We studied a free-ranging population of rhesus macaques living on Cayo Santiago, a 15.2 ha island off the southeastern coast of Puerto Rico. The current population of ∼1700 individually recognized rhesus macaques living in 12 social groups is descended from 409 monkeys that were transported from India to the island in 1938. This population is managed by the Caribbean Primate Research Center (CPRC) of the University of Puerto Rico. Monkeys are fed monkey chow daily, and water catchments provide ad libitum access to drinking water. During the study period (1960–2022), observers monitored and recorded demographic events daily. These records included births, deaths, sex, maternal identification, sires when genetic data were available, and group emigration/immigration events. The island is free of predators and, as approved by the Institutional Animal Care and Use Committee (IACUC), there is no regular veterinary intervention. Daily total rainfall and mean maximum temperature data were obtained from the NOAA station in Rio Piedras, Puerto Rico. Over a 63-year period (1959– 2022), data was not recorded by this NOAA station for 21% of days. We imputed missing rainfall and temperature data using the ‘mice’ package (v. 3.14.0) in R (Buuren & Groothuis-Oudshoorn, 2011). This study included 659 individuals over 6 years of age, including 346 females and 313 males.

### Behaviors

Behavioral data were collected continuously on handheld computers. From 2010 to 2022, 17 observers conducted focal samples on adults in 6 social groups (groups F, HH, KK, R, S, and V). For groups HH in 2016, KK in 2017, and S in 2019, 5-minute focal animal samples were collected. For all other group-years, 10-minute focal animal samples were used. During focal samples, observers recorded activity state (e.g., resting, traveling, feeding) and social interactions with other adults. For social interactions, observers recorded type of behavior (e.g., vocal grunts, approaches, contact aggression, displacements), the identity of the partner, and whether the focal or partner initiated the interaction. Focal animals were selected randomly and data were balanced such that animals were sampled roughly equally across times of day and across the study period. Animals included in our analyses were focal followed for a mean of 4.5 hours per study year (range: 0.75-8.8 hours). State behaviors include grooming given and grooming received. Event behaviors include aggression (contact and noncontact aggression, threats, displacements), submission (avoids, fear grimaces, submits), self-directed behaviors (scratches, self-grooming), and vigilance (which we began measuring in 2013). We established dominance ranks for all adults of a given sex in a given year using the direction and outcome of submissive, win-loss, interactions, and we use continuous rank values (Brent et al., 2013).

### Early life adversity

We used historical demographic records to assess individual exposure to early life adversity. We considered 8 forms of potential early life adversity. Adversities are experiences hypothesized to lead an individual to allocate their resources in a suboptimal manner. Each form of adversity has been justified in our previous work (Patterson et al., 2024). To maximize our sample size, we exclude two variables that we previously used: maternal social connectedness and maternal rank at birth. To have included these variables, we would have removed all individuals born before 2010, when our focal behavioral data collection began. Here are the 8 adversity measures: **Maternal loss:** When an individual’s mother died (including natural death owing to causes such as disease or injury and permanent removal from the population) before the individual reached 4 years of age. This 4-year window includes the period during which young macaques are nutritionally dependent on their mothers, and the period during which young macaques are weaned but still socially dependent on their mothers. **Competing sibling:** When a sibling was born within 355 days of a subject, which represented the bottom quartile of interbirth intervals (IBIs) in our sample. Last-born offspring and individuals that died before their sibling was born could not experience this adversity. The presence of competing siblings was measured as a binary variable. **Group size:** We used group size as a proxy for within-group competition. Demographic records were used to construct group composition over the study period. Group size was defined as the number of adults (≥4 years of age) of both sexes in an individual’s social group on the day that individual was born and it was included in our models as a continuous variable. **Primiparity:** Being born to a first-time mother might limit the energetic resources available for the developing individual and how those resources are allocated. We used a binary measurement for primiparity: first born or not first born. **Kin network:** We used kin network size as a proxy for social support. We measured an individual’s maternal kin network size at birth as the number of living females over 4 years of age with a relatedness coefficient of at least 0.063. Kin network size was included as a continuous variable. **Rainfall:** We used total rainfall (i.e., a continuous variable) across the first year of life. **Temperature:** We averaged mean maximum daily temperatures (i.e., a continuous variable) across the first year of life**. Hurricane:** We recorded individual exposure to any of the three major hurricanes (Category ≥3) that had major impacts on Cayo Santiago (Hugo on 18 September 1989, Georges on 21 September 1998, and Maria on 20 September 2017) during the first year of life. We treated this as a categorical variable to assess the effects of each major storm. **Cumulative early life adversity index:** the 8 adversity measures were summed together to create an index of cumulative exposure to early life adversity for each individual in our sample. Each adversity measure ranged between 0 and 1. Binary variables were scored as 0 (no adversity) or 1 (adversity) and continuous variables were adjusted to range from 0 to 1 continuously.

### Data analysis

The outcome variables were our behaviors of interest: self-directed behavior, vigilance, aggression given, aggression received, submissions, proportion of time grooming others, proportion of time being groomed, approaches given, and approaches received. For each year an individual was observed, we calculated the total counts of each behavior, the total duration of grooming, and the total number of hours an individual was observed. Grooming was represented as the proportion of observation time spent grooming others or being groomed. The other behaviors were represented as counts. Early life adversity was our main predictor variable of interest. Early life adversity variables were modelled in two ways: (i) cumulative index model, which included all forms of adversity summed together into one variable; and (ii) multivariate model, which included all eight forms of adversity, modelled as separate predictor variables in the same model. We assessed the efficacy of these two approaches by comparing the difference in the expected predictive accuracy for the cumulative index models versus the multivariate models. Final models were run two ways. Model 1 was our main model which included early life adversity index (or all individual adversity variables), current adult rank, sex, and age. Model 2 included additional interaction terms: an interaction term between early life adversity and adult rank, and between early life adversity and sex. Models also included a varying intercept for animal ID, social group ID, and year of behavioral data collection, and in the case of count data (i.e., all response variables except proportion of time grooming), an offset term for focal observation time.

Analyses were run in R (v. 4.1.2 (RStudio Team, 2021)) and RStudio (v. 1.4.1106 (R Core Team, 2021)). Models were run with the brms package (v. 2.16.3) (Bürkner, 2018). For count data, we specified a negative binomial error structure, and for proportion data, we specified a zero-inflated beta error structure. All continuous predictor variables were standardized to a mean of 0 and a standard deviation of 1. All models were Bayesian and we used weakly informative priors for fixed effects, setting the mean to zero and the standard deviation to one. Given the interrelatedness among behavioral outcome variables, we employed a multivariate response model (i.e., multiple outcome variables) for model 1 (main effects model) and a multivariate response model for model 2 (interactions model). Multivariate models specified that the outcome variables were predicted by a shared, correlated random intercept for each animal ID, group ID, and year of observation. This approach allows for borrowing strength across outcomes and provides more accurate estimates of uncertainty. We used credible intervals to determine whether the effect of a variable was substantial or not. If the 85% credible interval for an effect did not overlap with zero, the effect was considered substantial. When the vast majority of the 85% credible interval did not span zero, but there was some overlap, we described the model as being ‘uncertain’ about the effect. To compare how the cumulative index and multivariate models fit the data, we used the ‘loo’ model fit criterion in the brms package. Model code is available: https://github.com/skpatter/ELA_Behavior_Macaques

## Results

### Model 1: Main effects

Cumulative early life adversity predicted aspects of adult behavior (Table 1). Macaques who experienced more early life adversity received more aggression than those with less adversity (β = 0.03 ± 0.03 (mean of the posterior distribution ± the posterior standard deviation); Fig 1). Early life adversity was associated with more submissiveness (β = 0.05 ± 0.03) and less vigilance (β = −0.01 ± 0.01; Fig 1). Individuals with more early life adversity spent less time being groomed by others compared to individuals with less early life adversity (β = −0.05 ± 0.03; Fig 1). There were no observable effects of early life adversity on aggressiveness, self-directed behaviors, time spent grooming others, or approaches (Fig 1). Full model results are presented in supplementary materials (SOM Table S1). Models with the cumulative early life adversity index fit the data better than models with each form of early life adversity included as separate predictor variables (SOM Table S3).

**Figure 1.**
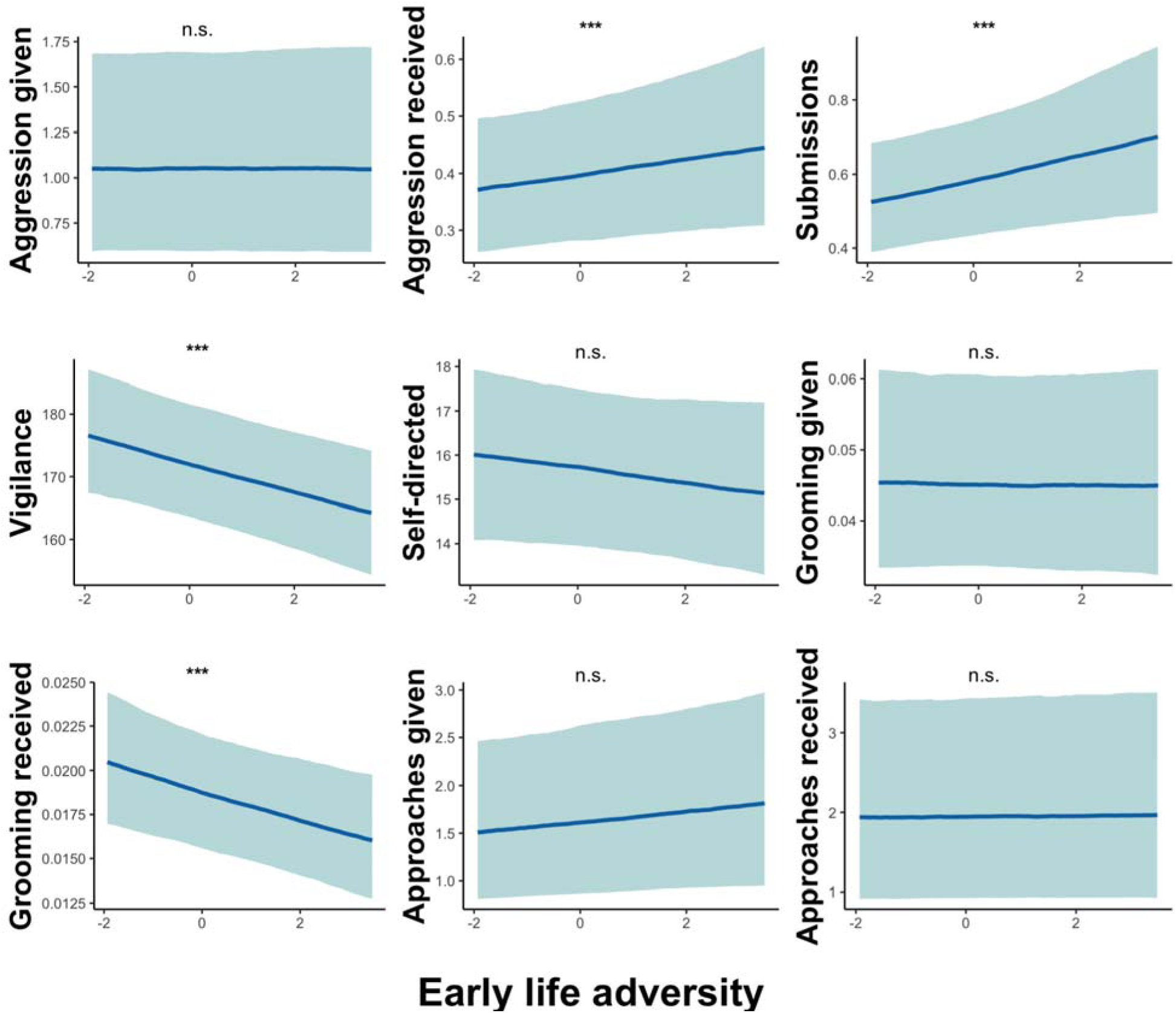
Model predictions are shown for the effect of cumulative early life adversity on behavior in adulthood. Predictions are from model 1 and demonstrate the main effects of early life adversity while holding other model covariates constant. Early life adversity on the x-axis is standardized such that zero represents the mean amount of early life adversity experienced in the sample. Three asterisks (***) denote substantial effects and “n.s.” denotes non-substantial effects for which the credible intervals overlapped with zero.

**Table 1.** Summary of behaviors, predictions, and findings.

| Behavior Type | Behavior | Prediction | Support? |
| --- | --- | --- | --- |
| Self-directed | Self-scratching, self-grooming | More ELA -> higher rates | No effect observed |
| Vigilance | Watchfulness or attentiveness to surroundings | More ELA -> higher rates | Mixed; rank-dependent |
| Agonistic | Aggression (given, received)<br>Submission | More ELA -> higher rates | Mixed; rank-dependent for aggression received and submissions |
| Affiliative | Grooming (given, received)<br>Approaches (given, received) | More ELA -> lower rates | Mixed; rank- and sex-dependent |

### Model 2: Interaction effects

Some of the effects of cumulative early life adversity on behavior were modulated by sex and rank (Table 1). There was an interaction effect between early life adversity and sex on vigilance and approaches received. The effect of adversity on vigilance was stronger for males than females with males exhibiting a steeper decline in vigilance with greater early life adversity (β = −0.04 ± 0.01; Fig 2). Females with more early life adversity received fewer approaches from others and males with more adversity received more approaches from others (β = 0.15 ± 0.05; Fig 2). There were not observable interaction effects between sex and early life adversity for the other behavioral outcomes (SOM Table S2).

**Figure 2.**
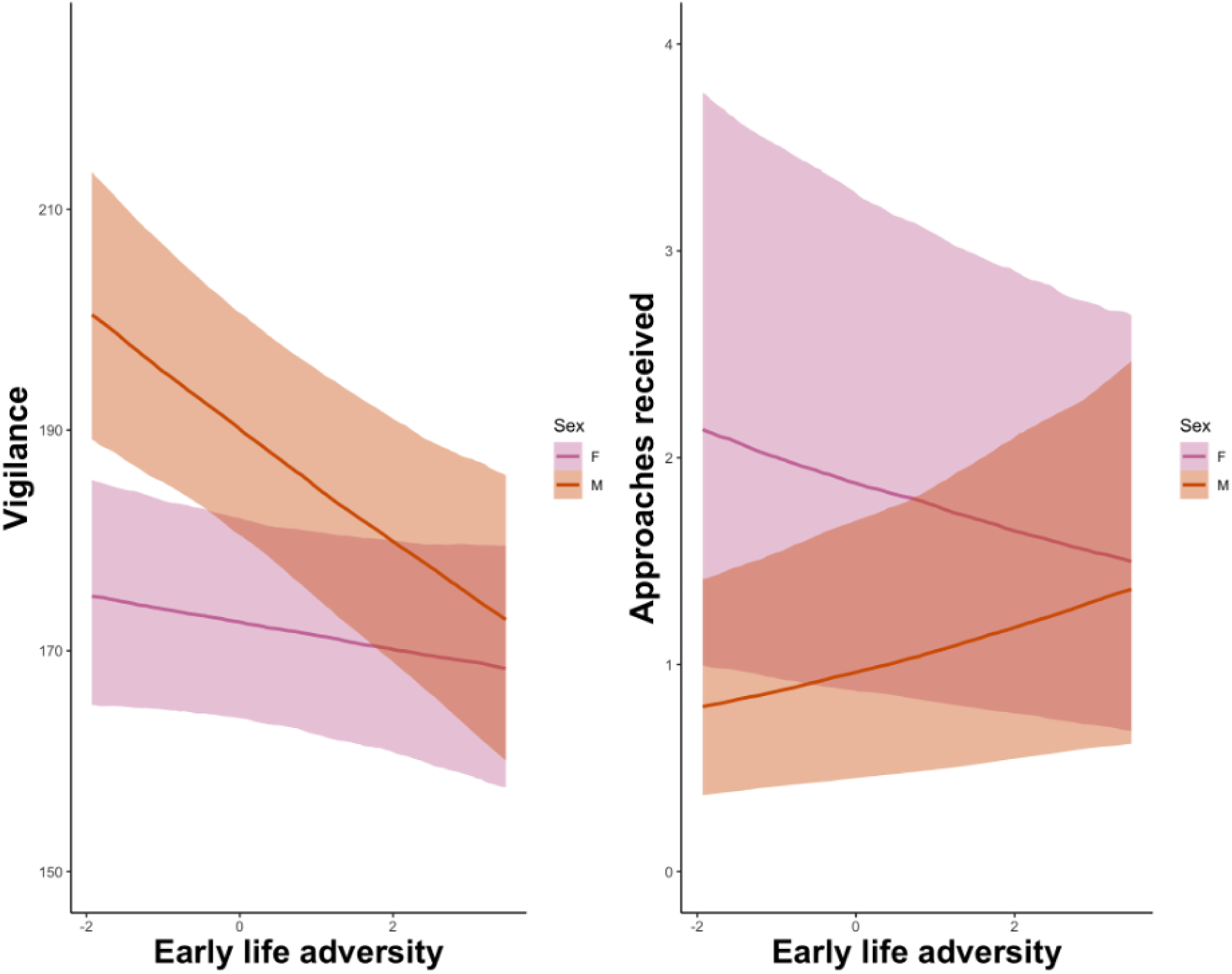
Model 2 predictions are shown for the interactive effects between cumulative early life adversity and sex on behavior. Early life adversity on the x-axis is standardized such that zero represents the mean amount of early life adversity experienced in the sample. Pink represents females and orange represents males.

Adult dominance rank moderated the effects of early life adversity on submissions, vigilance, aggression received, grooming received, and grooming given. Early life adversity was associated with more submissiveness and vigilance for higher ranking individuals but less submissiveness and vigilance for lower ranking individuals (submissions: β = −0.08 ± 0.03; vigilance: β = −0.02 ± 0.01; Fig 3). The positive association between early life adversity and aggression received was strongest for high ranking individuals and weakest for lower ranking individuals (β = −0.06 ± 0.03; Fig 3). More early life adversity was associated with less grooming given and received for low ranking individuals, while for high ranking individuals, early life adversity had no observable effect on grooming received and a positive effect on grooming given (Given: β = −0.05 ± 0.03; received: β = −0.06 ± 0.03; Fig 3). There were no observable interaction effects between rank and early life adversity for the other behavioral outcomes (SOM Table S2).

**Figure 3.**
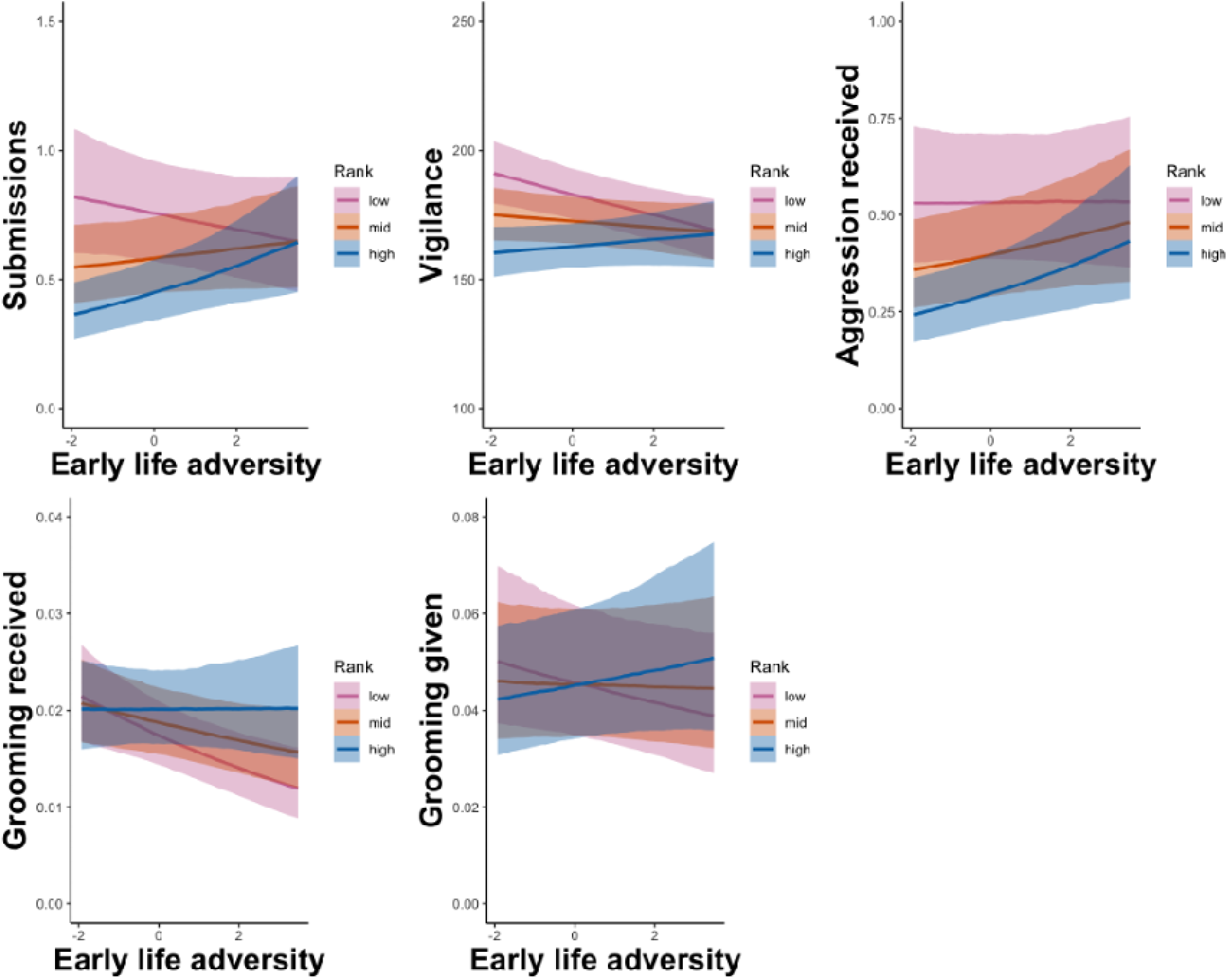
Model 2 predictions are shown for the interactive effects between cumulative early life adversity and adult rank on behavior. Early life adversity on the x-axis is standardized such that zero represents the mean amount of early life adversity experienced in the sample. Red represents low rank, orange represents mid rank, and blue represents high rank.

## Discussion

By integrating long-term demographic and ecological data with detailed focal behavioral data, we were able to assess how early life adversity predicts adult behavior in a free-ranging population of rhesus macaques. We found that cumulative early life adversity shaped the ways in which macaques interacted with conspecifics and their environment. Individuals that experienced more early life adversity received more aggression and less grooming from others, were more submissive, and were less vigilant. Some effects were modulated by rank and sex, illustrating the complexities of early life effects and behavioral strategies.

Although we predicted that individuals with early life adversity would develop more aggressive phenotypes and as a result, receive more aggression, we only found partial support for this prediction. Early life adversity did not appear to shape how aggressive an individual was in adulthood. The lack of evidence for increased aggression is noteworthy because early life adversity has been associated with patterns of aggression in primates, including humans, and in rodents (Condon et al., 2022; Haller et al., 2014; Suomi, 1997; Veenema, 2009). However, field crickets show reduced aggression following cues of high competition (DiRienzo et al., 2012), and while rodents typically increase aggression following maternal separation, some studies have found that male mice decrease aggression (Veenema, 2009). Additionally, dogs exposed to early life adversity show higher levels of aggression and fear as adults, but the strength of effect varies by breed suggesting genetic differences in sensitivity (Espinosa et al., 2025). While early life adversity often leads to increased aggression, this behavioral response seems to be context-dependent, varying by species, sex, genes (e.g., dog breed), and socioecology (e.g., captive versus non-captive environments; form or context of adversity).

We found that early life adversity predicted the amount of aggression macaques received. Individuals that experienced more early life adversity were more likely to be targets of aggression than those with less early life adversity. Contrary to our predictions, rather than being buffered against this consequence, high ranking individuals appeared to receive more aggression as a function of adversity than lower ranking individuals. Receiving frequent aggression from others could be one factor shaping mortality risk via higher rates of wounds, infection, and disease (Cooper et al., 2025; Patterson et al., 2024; Pavez-Fox et al., 2022, 2025). One explanation for this finding could be that those who experienced early life adversity were easier physical targets, perhaps due to small body size, disease, or limited motor skills, and thus were targeted more. If this were the sole explanatory factor, it is not clear why high ranking individuals would be most affected. Other aspects of social behavior could also shape why these individuals were targeted. In other species, early life adversity has been linked to predictability, boldness, and social competence or the ability to respond appropriately in various social contexts (Heiming et al., 2009; Patterson et al., 2022; Taborsky et al., 2012). Rhesus macaques exposed to early life adversity might employ different social phenotypes based on their social rank, which could affect the amount of aggression others direct towards them.

Early life adversity predicted affiliative interactions, and the nature of some of these effects varied as a function of rank and sex. Greater early life adversity was associated with less time grooming, a pattern in line with previous work on female baboons (Lange et al., 2023; Patterson et al., 2022; Rosenbaum et al., 2020). Among lower ranking individuals in particular, those with more early life adversity spent less time grooming and being groomed. Higher ranking individuals seem to have been buffered against these consequences. With regard to spatial proximity, different phenotypes emerged for males and females. Females with more early life adversity were approached less than those with less adversity; while males with more early life adversity were approached more than those with less adversity. The higher rates of approaches for males with early life adversity is unexpected. Being frequently approached could be a signal that high adversity males are socially attractive to others, or that they are non-threatening individuals to approach. Alternatively, these approaches could be a precursor to aggression. Future analyses investigating the dyadic composition of interactions and quality of social bonds will be needed to shed light on these phenotypic outcomes.

Some but not all behavioral markers associated with stress and fear responses were shaped by early life experience. Patterns of submissive behavior mapped onto patterns of aggression received such that individuals with early life adversity were targeted more aggressively and were more likely to exhibit submissiveness compared to those without adversity. Furthermore, high ranking individuals, as opposed to lower ranking individuals, were targeted the most as a function of early life adversity, and were also more likely to exhibit greater submissiveness and vigilance in relation to greater exposure to early life adversity. On the other hand, among lower ranking individuals, early life adversity did not appear to result in substantially higher levels of aggression received but did predict lower levels of submissiveness and vigilance. This means early life adversity might not directly lead to higher levels of these stress-related behaviors, but rather individuals might be appropriately responding to their social environment. Early life adversity is associated with higher glucocorticoids and altered HPA function in humans, baboons, spotted hyenas, rodents, voles, and juvenile rhesus macaques in this study population (Hunter et al., 2011; Laubach et al., 2019; Patterson et al., 2021; Petrullo et al., 2016; Pisu et al., 2016; Rosenbaum et al., 2020; Wei et al., 2013; Young et al., 2019). While these physiological findings might suggest that macaques exposed to early life adversity struggle to cope with environmental challenges, further work is needed to assess whether these outcomes persist into adulthood.

Consistent with prior research on wild populations, our analysis indicates that cumulative early life adversity explained behavioral outcomes better than individual measures of adversity (Patterson et al., 2021, 2022; Tung et al., 2016). The accumulation of adversities appears more influential for behavioral development than any single form of adversity measured. However, this does not imply that all forms of adversity produce the same impact. In humans, distinct types of adversity, such as deprivation versus threats, are expected to produce divergent behavioral outcomes (McLaughlin & Sheridan, 2016; Sheridan et al., 2020). Future research is warranted to disentangle the effects of individual adversities, dimensions of adversity, and cumulative adversity.

Several lines of further inquiry may prove fruitful for understanding behavioral responses to early life adversity. First, assessing how variation in behavioral strategies correlates with fitness outcomes would help us identify why different behavioral strategies arise. Male and female rhesus macaques attain social status in different ways, navigate different competitive landscapes, and experience different social environments (Brent et al., 2017b; Cooper et al., 2024; Higham & Maestripieri, 2014; Hoffman et al., 2008). Behavioral outcomes like reduced social connectedness are detrimental for survival and reproduction on average (Snyder-Mackler et al., 2020), but could be beneficial for certain individuals depending on context. For example, isolating from others could be an effective aggression-avoidance strategy for individuals navigating the misfortune of a challenging developmental period and low adult social status. In contrast, high ranking individuals with early adversity may not benefit from isolating as they would miss out on social benefits of high status. In this population, extreme aggressive phenotypes – being very aggressive or minimally aggressive – yield the best reproductive output, while moderate levels of aggressiveness fare the worst (Brent et al., 2013). Similar analyses testing reproductive success conditional upon early life adversity and social status will help clarify the mechanisms and pressures driving behavioral variation. Second, identifying the developmental pathways underlying the link between early life adversity and adult behaviors may help elucidate the fitness consequences of behavioral responses to adversity. Factors such as sex and resource access can shape how individuals navigate developmental trade-offs (Berghänel et al., 2015; Patterson et al., 2025), and could ultimately lead to different adult phenotypes. Phenotypic adjustments in response to adversity are expected to provide immediate survival benefits, but might lead to detrimental outcomes later in life, or such adjustments could prove beneficial in adulthood as well (Hinde, 2013; Lea & Rosenbaum, 2020; Lu et al., 2019; Nettle et al., 2013; Nettle & Bateson, 2015; Patterson et al., 2022). Third, investigating who (e.g., group-mates, same sex, kin) is aggressing on individuals with early life adversity, which aspects of the targeted individual’s behavior or health explains the rate of aggression received, and whether aggression received mediates the link between early life adversity and mortality risks will be important avenues for future research in this population.

Our analysis revealed several general behavioral patterns associated with early life adversity, as well as nuanced patterns of early life adversity and behavior suggesting different strategies might be employed based on variables like social status and sex. Early life adversity predicted how others interacted aggressively and affiliatively with focal individuals. We found evidence for different effects of early life adversity for high versus low status individuals, which did not necessarily align with buffering effects. High social status appeared to mitigate the negative association between early life adversity and grooming, but amplify the positive association between adversity and aggression received. While individual sex shaped the relationship between early life adversity and two behaviors (proximity and vigilance), early adversity had largely similar effects on male and female behavior. The fitness correlates of behavioral phenotypes might vary according to factors like sex, rank, health, current environmental conditions, and kin support (Brent et al., 2013, 2017a; Testard et al., 2024).

Our study demonstrates how observing animals in non-captive environments enhances our understanding of early life sensitivities across species. While human studies are often limited by sociocultural confounding variables, animal models offer the ability to isolate effects of adversity at specific times. However, captive primate and rodent experiments rely on artificial adversities and unnatural socioecological environments. Examining naturally occurring adversity under natural conditions better reflects evolutionarily relevant selective pressures. For instance, unlike captive rhesus macaque studies, we found little evidence that macaques with early life adversity displayed higher rates of aggression or self-directed behaviors. This divergence may stem from experimental differences (e.g., testing behavior during an experimental challenge), the severity of adversities, or the lack of agency in captive groupings compared to natural conditions where individuals can freely range and manage relationships. Our analysis adds to previous work, which found that early life adversity in wild primates increases the risk of adult social isolation, by showing that adversity also shapes agonism, vigilance, and how group mates interact with an individual, both affiliatively and aggressively (Lange et al., 2023; Patterson et al., 2022; Rosenbaum et al., 2020; Tung et al., 2016). Expanding research into natural populations will better elucidate the mechanisms, strategies, and fitness correlates of responses to early life adversity, ultimately providing deeper insight into human behavior and health risks

## Data sharing

Data used for this manuscript are available here: https://zenodo.org/records/21419720 [10.5281/zenodo.21419720]

## Acknowledgments

We would like to thank the Caribbean Primate Research Center at the University of Puerto Rico, in particular the Cayo Santiago field team, for the long-term collection of the data presented here. This research was supported by funding from the National Science Foundation (SMA-2105307; BCS-1800558; BCS-1754024) and National Institutes of Health (R01-AG060931; R01-AG084706; R56-AG071023; R00-AG051764; R01-MH096875; R01-MH089484; R01-MH118203). The Cayo Santiago field station is supported by the University of Puerto Rico and the Office of Research Infrastructure Programs of the NIH grant number P40-OD012217.

