## Supplementary Materials for "Early life adversity shapes adult behavior in a free-ranging primate"

**Table S1. Model 1 output: Main effects**

| | Estimate | Est.Error | l-85% CI | u-85% CI |
|---|---|---|---|---|
| apprec_Intercept | 0.64 | 0.48 | -0.07 | 1.24 |
| appgive_Intercept | 0.46 | 0.4 | -0.12 | 0.98 |
| subgive_Intercept | -0.58 | 0.2 | -0.86 | -0.31 |
| agggive_Intercept | 0.04 | 0.37 | -0.51 | 0.53 |
| aggrec_Intercept | -0.97 | 0.24 | -1.3 | -0.67 |
| vigcount_Intercept | 5.14 | 0.04 | 5.09 | 5.2 |
| selfcount_Intercept | 2.75 | 0.08 | 2.63 | 2.86 |
| AdultGroomGiveProp_Intercept | -2.9 | 0.22 | -3.2 | -2.58 |
| AdultGroomGetProp_Intercept | -3.76 | 0.13 | -3.95 | -3.59 |
| apprec_s_age | 0.03 | 0.02 | -0.01 | 0.06 |
| apprec_s_CumulIndex | 0 | 0.03 | -0.04 | 0.04 |
| apprec_s_rank | -0.08 | 0.03 | -0.12 | -0.04 |
| apprec_sexM | -0.67 | 0.05 | -0.74 | -0.59 |
| appgive_s_age | -0.08 | 0.03 | -0.12 | -0.05 |
| appgive_s_CumulIndex | 0.03 | 0.03 | -0.01 | 0.07 |

| | | | | |
|---|---|---|---|---|
| appgive_s_rank | -0.2 | 0.03 | -0.24 | -0.15 |
| appgive_sexM | -0.52 | 0.05 | -0.6 | -0.44 |
| subgive_s_age | -0.01 | 0.03 | -0.05 | 0.03 |
| subgive_s_CumulIndex | 0.05 | 0.03 | 0.01 | 0.1 |
| subgive_s_rank | 0.26 | 0.03 | 0.22 | 0.31 |
| subgive_sexM | -0.5 | 0.06 | -0.59 | -0.41 |
| agggive_s_age | -0.03 | 0.02 | -0.07 | 0.01 |
| agggive_s_CumulIndex | 0 | 0.02 | -0.04 | 0.04 |
| agggive_s_rank | -0.1 | 0.03 | -0.14 | -0.06 |
| agggive_sexM | 0.39 | 0.06 | 0.31 | 0.47 |
| aggrec_s_age | -0.28 | 0.03 | -0.33 | -0.24 |
| aggrec_s_CumulIndex | 0.03 | 0.03 | 0 | 0.08 |
| aggrec_s_rank | 0.3 | 0.03 | 0.26 | 0.34 |
| aggrec_sexM | -0.12 | 0.06 | -0.21 | -0.03 |
| vigcount_s_age | -0.05 | 0.01 | -0.06 | -0.04 |
| vigcount_s_CumulIndex | -0.01 | 0.01 | -0.02 | 0 |
| vigcount_s_rank | 0.06 | 0.01 | 0.05 | 0.07 |
| vigcount_sexM | 0.1 | 0.01 | 0.08 | 0.12 |
| selfcount_s_age | -0.06 | 0.01 | -0.08 | -0.04 |
| selfcount_s_CumulIndex | -0.01 | 0.01 | -0.03 | 0.01 |
| selfcount_s_rank | 0.04 | 0.01 | 0.02 | 0.06 |
| selfcount_sexM | 0.22 | 0.02 | 0.19 | 0.26 |
| AdultGroomGiveProp_s_age | -0.04 | 0.03 | -0.08 | -0.01 |
| AdultGroomGiveProp_s_CumulIndex | 0 | 0.03 | -0.05 | 0.04 |

| | | | | |
|---|---|---|---|---|
| AdultGroomGiveProp_s_rank | 0.01 | 0.03 | -0.04 | 0.05 |
| AdultGroomGiveProp_sexM | -0.26 | 0.06 | -0.35 | -0.17 |
| AdultGroomGetProp_s_age | 0.02 | 0.03 | -0.02 | 0.05 |
| AdultGroomGetProp_s_CumulIndex | -0.05 | 0.03 | -0.09 | -0.01 |
| AdultGroomGetProp_s_rank | -0.08 | 0.03 | -0.12 | -0.04 |
| AdultGroomGetProp_sexM | 0.07 | 0.06 | -0.01 | 0.15 |

**Table S2. Model 2 output: Interaction effects**

| | Estimate | Est.Error | l-85% CI | u-85% CI |
|---|---|---|---|---|
| apprec_Intercept | 0.6 | 0.48 | -0.13 | 1.19 |
| appgive_Intercept | 0.44 | 0.41 | -0.13 | 0.97 |
| subgive_Intercept | -0.58 | 0.19 | -0.83 | -0.32 |
| agggive_Intercept | 0.01 | 0.37 | -0.52 | 0.5 |
| aggrec_Intercept | -0.96 | 0.22 | -1.27 | -0.66 |
| vigcount_Intercept | 5.14 | 0.04 | 5.09 | 5.2 |
| selfcount_Intercept | 2.75 | 0.08 | 2.63 | 2.86 |
| AdultGroomGiveProp_Intercept | -2.88 | 0.22 | -3.17 | -2.57 |
| AdultGroomGetProp_Intercept | -3.77 | 0.14 | -3.95 | -3.58 |
| apprec_s_age | 0.02 | 0.02 | -0.01 | 0.06 |
| apprec_s_CumulIndex | -0.06 | 0.03 | -0.11 | -0.02 |
| apprec_s_rank | -0.08 | 0.03 | -0.12 | -0.04 |
| apprec_sexM | -0.67 | 0.05 | -0.75 | -0.59 |
| apprec_s_CumulIndex:s_rank | -0.01 | 0.03 | -0.05 | 0.02 |
| apprec_s_CumulIndex:sexM | 0.16 | 0.05 | 0.09 | 0.23 |

| | | | | |
|---|---|---|---|---|
| appgive_s_age | -0.08 | 0.03 | -0.12 | -0.04 |
| appgive_s_CumulIndex | 0.01 | 0.03 | -0.04 | 0.06 |
| appgive_s_rank | -0.2 | 0.03 | -0.25 | -0.16 |
| appgive_sexM | -0.52 | 0.06 | -0.6 | -0.44 |
| appgive_s_CumulIndex:s_rank | 0.03 | 0.03 | -0.01 | 0.07 |
| appgive_s_CumulIndex:sexM | 0.05 | 0.05 | -0.02 | 0.12 |
| subgive_s_age | -0.02 | 0.03 | -0.06 | 0.02 |
| subgive_s_CumulIndex | 0.04 | 0.04 | -0.01 | 0.09 |
| subgive_s_rank | 0.28 | 0.03 | 0.23 | 0.32 |
| subgive_sexM | -0.51 | 0.06 | -0.6 | -0.42 |
| subgive_s_CumulIndex:s_rank | -0.08 | 0.03 | -0.12 | -0.04 |
| subgive_s_CumulIndex:sexM | 0.06 | 0.06 | -0.02 | 0.14 |
| agggive_s_age | -0.03 | 0.02 | -0.06 | 0 |
| agggive_s_CumulIndex | 0 | 0.03 | -0.05 | 0.05 |
| agggive_s_rank | -0.1 | 0.03 | -0.14 | -0.06 |
| agggive_sexM | 0.39 | 0.05 | 0.32 | 0.47 |
| agggive_s_CumulIndex:s_rank | -0.01 | 0.03 | -0.05 | 0.03 |
| agggive_s_CumulIndex:sexM | 0 | 0.05 | -0.07 | 0.06 |
| aggrec_s_age | -0.29 | 0.03 | -0.33 | -0.25 |
| aggrec_s_CumulIndex | 0.06 | 0.04 | 0.01 | 0.11 |
| aggrec_s_rank | 0.31 | 0.03 | 0.26 | 0.35 |
| aggrec_sexM | -0.12 | 0.06 | -0.21 | -0.03 |
| aggrec_s_CumulIndex:s_rank | -0.06 | 0.03 | -0.1 | -0.01 |
| aggrec_s_CumulIndex:sexM | -0.05 | 0.06 | -0.13 | 0.03 |

| | | | | |
|---|---|---|---|---|
| vigcount_s_age | -0.05 | 0.01 | -0.06 | -0.05 |
| vigcount_s_CumulIndex | -0.01 | 0.01 | -0.02 | 0.01 |
| vigcount_s_rank | 0.06 | 0.01 | 0.05 | 0.07 |
| vigcount_sexM | 0.1 | 0.01 | 0.08 | 0.12 |
| vigcount_s_CumulIndex:s_rank | -0.02 | 0.01 | -0.03 | -0.01 |
| vigcount_s_CumulIndex:sexM | -0.02 | 0.01 | -0.04 | 0 |
| selfcount_s_age | -0.06 | 0.01 | -0.08 | -0.04 |
| selfcount_s_CumulIndex | -0.01 | 0.02 | -0.03 | 0.01 |
| selfcount_s_rank | 0.04 | 0.01 | 0.02 | 0.06 |
| selfcount_sexM | 0.22 | 0.02 | 0.19 | 0.26 |
| selfcount_s_CumulIndex:s_rank | -0.01 | 0.01 | -0.03 | 0.01 |
| selfcount_s_CumulIndex:sexM | 0 | 0.02 | -0.04 | 0.03 |
| AdultGroomGiveProp_s_age | -0.05 | 0.03 | -0.09 | -0.01 |
| AdultGroomGiveProp_s_CumulIndex | 0 | 0.04 | -0.05 | 0.05 |
| AdultGroomGiveProp_s_rank | 0.01 | 0.03 | -0.04 | 0.06 |
| AdultGroomGiveProp_sexM | -0.27 | 0.06 | -0.36 | -0.18 |
| AdultGroomGiveProp_s_CumulIndex:s_rank | -0.05 | 0.03 | -0.09 | 0 |
| AdultGroomGiveProp_s_CumulIndex:sexM | -0.02 | 0.06 | -0.1 | 0.07 |
| AdultGroomGetProp_s_age | 0.01 | 0.03 | -0.03 | 0.05 |
| AdultGroomGetProp_s_CumulIndex | -0.05 | 0.03 | -0.1 | 0 |
| AdultGroomGetProp_s_rank | -0.08 | 0.03 | -0.12 | -0.03 |
| AdultGroomGetProp_sexM | 0.07 | 0.06 | -0.02 | 0.16 |
| AdultGroomGetProp_s_CumulIndex:s_rank | -0.06 | 0.03 | -0.1 | -0.01 |
| AdultGroomGetProp_s_CumulIndex:sexM | -0.01 | 0.06 | -0.09 | 0.07 |

**Table S3.** WAIC model comparison for Cumulative ELA and Separate ELA variables

| | | |
|---|---|---|
| **Index is better** for self-directed behaviors**:** | | |
| | elpd_diff | se_diff |
| Self-directed Cumulative Index | 0.0 | 0.0 |
| Self-directed Separate | -4.1 | 1.9 |
| **Index is better** for vigilance**:** | | |
| | elpd_diff | se_diff |
| Vigilance Cumulative Index | 0.0 | 0.0 |
| Vigilance Separate | -5.5 | 2.1 |
| No clear difference for aggression given: | | |
| | elpd_diff | se_diff |
| Aggression Give Cumulative Index | 0.0 | 0.0 |
| Aggression Give Separate | -1.1 | 2.4 |
| No clear difference for aggression received: | | |
| | elpd_diff | se_diff |
| Aggression Receive Separate | 0.0 | 0.0 |
| Aggression Received Cumulative Index | -3.8 | 4.5 |
| No clear difference for submissions: | | |
| | elpd_diff | se_diff |
| Submission Cumulative Index | 0.0 | 0.0 |
| Submission Separate | -0.8 | 3.1 |

| | | |
|---|---|---|
| **Index is better** for grooming given**:** | | |
| | elpd_diff | se_diff |
| Groom Give Cumulative Index | 0.0 | 0.0 |
| Groom Give Separate | -4.4 | 2.6 |
| No clear difference for grooming received: | | |
| | elpd_diff | se_diff |
| Groom Get Separate | 0.0 | 0.0 |
| Groom Get Cumulative Index | -0.7 | 3.7 |
| **Index is better** for approaches given to others**:** | | |
| | elpd_diff | se_diff |
| Approach Give Cumulative Index | 0.0 | 0.0 |
| Approach Give Separate | -4.4 | 3.3 |
| No clear difference for approaches received: | | |
| | elpd_diff | se_diff |
| Approach Receive Cumulative Index | 0.0 | 0.0 |
| Approach Receive Separate | -1.8 | 3.5 |
